# Centromere Pairing in Prophase Allows Partner Chromosomes to Orient on the Meiosis I Spindle

**DOI:** 10.1101/2022.12.19.520819

**Authors:** Jared M. Evatt, Hoa H. Chuong, Régis E. Meyer, Dean S. Dawson

**Affiliations:** Oklahoma Medical Research Foundation, Oklahoma City, Oklahoma, USA, 73104; Department of Cell Biology, University of Oklahoma Health Sciences Center, Oklahoma City, Oklahoma, USA

## Abstract

Proper chromosome segregation in meiosis I relies on the formation of connections between homologous chromosomes. Crossovers between homologs provide a connection that allows them to attach correctly to the meiosis I spindle. Tension is transmitted across the crossover when the partners attach to microtubules from opposing poles of the spindle. Tension stabilizes microtubule attachments that will pull the partners towards opposite poles at anaphase ^1,2^. Paradoxically, in many organisms, non-crossover partners segregate correctly ^3^. The mechanism by which non-crossover partners become bi-oriented on the meiotic spindle is unknown. Both crossover and noncrossover partners pair their centromeres in early in meiosis (prophase). In budding yeast, centromere pairing, is correlated with subsequent correct segregation of the partners ^4,5^. The mechanism by which centromere pairing, in prophase, promotes later correct attachment of the partners to the metaphase spindle is unknown. We used live cell imaging to track the bi-orientation process of non-crossover chromosomes. We find that centromere pairing allows the establishment of connections between the partners that allows their later interdependent attachment to the meiotic spindle using tension-sensing bi-orientation machinery. Because all chromosome pairs experience centromere pairing, our findings suggest that crossover chromosomes also utilize this mechanism to achieve maximal segregation fidelity.

---

Improper segregation of homologous chromosomes in meiosis I results in the production of aneuploid gametes which are a major cause of miscarriage, infertility, and birth defects^6^. Proper segregation in meiosis I relies upon the formation of physical linkages that connect homologous partner chromosomes to one another. The existing paradigm suggests that genetic recombination via crossing-over (or exchanges) results in the formation of chiasmata, which along with sister chromatid cohesion distal to the crossovers, serve as the basis of interhomolog linkages (reviewed in^7^). Chiasmata allow for tension to be applied across the homologous chromosome superstructure, called a bivalent, when the homologous partners are connected to opposite poles of the meiotic spindle (called bi-orientation). This tension at the kinetochore-microtubule interfaces of bi-oriented partners stabilizes the microtubule connections^1,2,8^. Homologous partners that fail to form crossovers (non-exchange partners) are more prone to segregation errors than exchange partners. However, studies in model organisms suggest that these non-exchange chromosomes do not segregate randomly in meiosis I^9–13^.

In recent years, the discovery of centromere pairing in meiotic prophase in multiple organisms has raised interesting questions about how this process may be positively affecting the segregation fidelity of both exchange and non-exchange homologous partners in meiotic anaphase I^4,5,14–16^. In yeast, Drosophila females and mouse spermatocytes, homologous centromeres become paired or clustered (in *Drosophila* females) by components of the synaptonemal complex (SC)^4,17^. This centromere pairing of non-exchange partners in prophase has been shown in yeast to be correlated with their subsequent disjunction in meiosis I^4,5,16^. A separation-of-function allele of the synaptonemal complex gene *ZIP1, zip1-N1*, which ablates centromere pairing, randomizes the segregation of non-exchange partners in anaphase I^18^.

Here, we used live cell imaging approaches to address how pairing of partner centromeres in prophase could impact attachment to the spindle and subsequent segregation at anaphase I.

## Prophase centromere pairing dissolves in pro-metaphase

One model that explains the role of centromere pairing in mediating the correct segregation of partner chromosomes is that the pairing persists through prometaphase, providing a tension-transmitting bridge that facilitates bi-orientation of the paired centromeres in pro-metaphase. A prediction of this model is that the centromeres would remain tethered through the bi-orientation process. To test this, we used live cell imaging to monitor the pro-metaphase behavior of centromeres that were paired in prophase. For these experiments we used well characterized mini-chromosomes (circular centromere plasmids) that act as partners in meiosis I ^19^. These mini-chromosomes carry natural yeast centromeres, origins of DNA replication, and do not become connected by crossovers. One mini-chromosome carries an array of *tet* operator repeats that are bound by a fluorescent mTurquoise-tetR fusion protein while the other carries a *lac* operator array and is bound by a yEVenus-lacI fusion protein. The cells also express a fluorescent red protein that localizes to the yeast microtubule organizing center (spindle pole body; SPB) and enables the determination of transitions from prophase to pro-metaphase and metaphase to anaphase. Live cell imaging analysis was performed by first allowing cultures of meiotic cells to arrest in the pachytene stage of prophase I then collecting images (every two minutes) of the cells in microfluidics chambers following release from the pachytene arrest. The pro-metaphase behavior of prophase paired centromeres was tracked in nineteen cells (Fig. 1). The average duration of prometaphase (from spindle formation to spindle elongation) was 79 minutes. In all nineteen cells the centromere pair separated within six minutes of spindle formation and moved on the spindle as separated foci. Thus, centromere pairing does not persist as a direct link that promotes bi-orientation.

**Figure 1.**
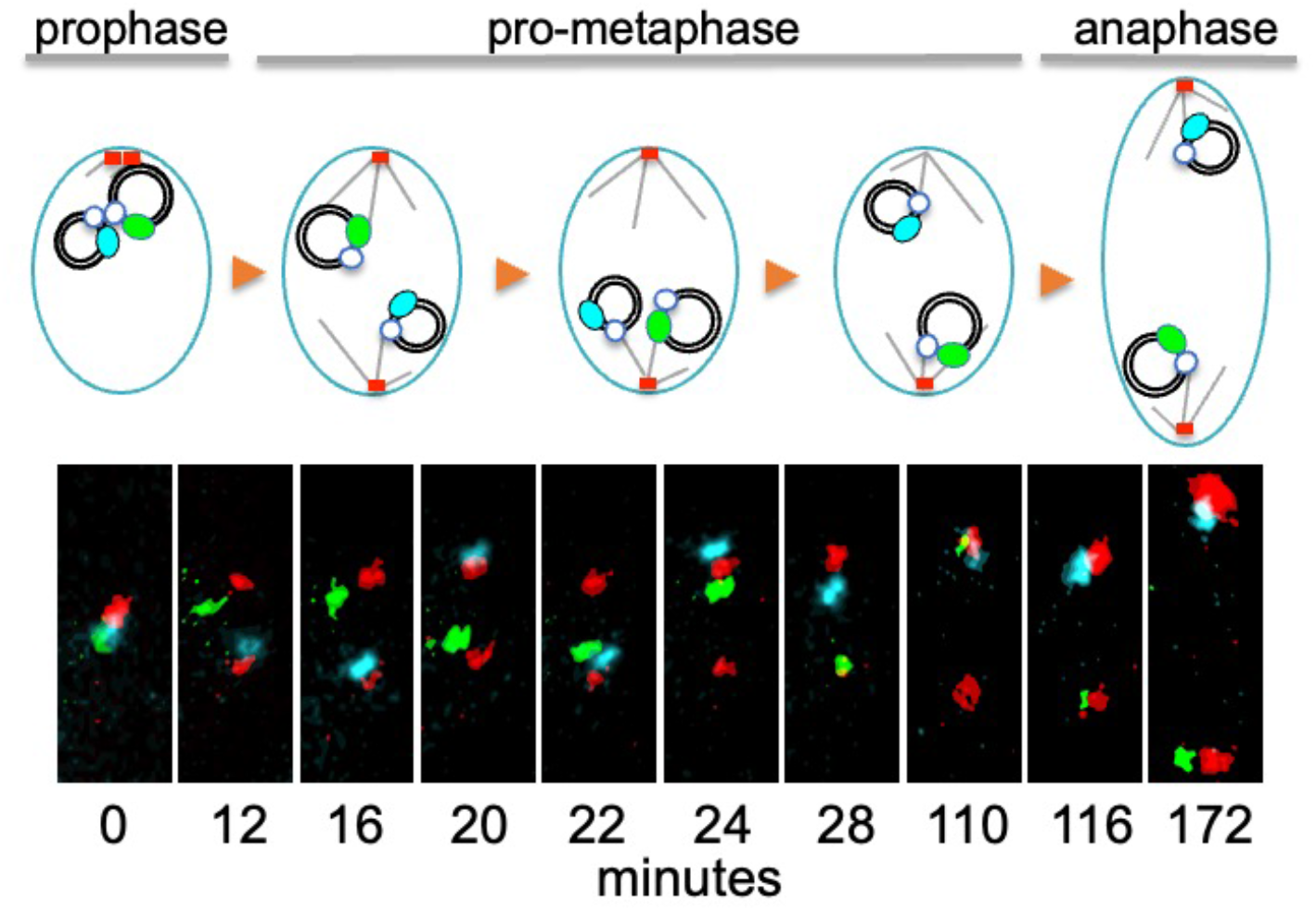
Paired centromeres dissociate upon entry into prometaphase. Live cell imaging was used to follow the fate in prometaphase of centromeres that were paired in meiotic prophase. Spindle pole bodies are marked in red (Spc42-DSRed) and mini-chromosome partners are marked by mTurquoise (blue) and mVenus (green). Images were acquired every two minutes. Selected images are shown. T=0 is the last frame before the SPBs separated, indicating spindle formation and the beginning of prometaphase. Anaphase entry is indicated by an inflection in spindle growth.

## Centromere pairing in prophase promotes inter-dependent centromere movements during the bi-orientation process

An alternate model to explain the impact of centromere pairing is that the tight juxtaposition it provides allows the chromatin of the partner chromosomes to be linked, and this link is what allows the centromeres to bi-orient in pro-metaphase. Such a connection is not easily visualized in our imaging experiments. However, the model predicts the pair will exhibit interdependent movements in prometaphase. To test this, we measured the distance between the minichromosome centromeres in the first of prometaphase in wild-type cells and in cells *(zip1-N1* mutants) that have reduced centromere pairing. As controls, we measured the movements of tagged, homologous chromosome centromeres (*CEN1*), connected by crossovers on the adjacent chromosome arms, or heterologous centromeres (one copy of *CEN1* and *CEN4*) each with their own homologous partner (Fig. 2A). We measured the fluctuations in inter-centromeric distance as each fluorescently tagged centromere pair moved on the spindle (Supl. Fig. 1 A). A mean-squared displacement analysis, revealed that the homologous partners (WT *CEN1/CEN1*) show low variation in centromere-centromere distance compared to the heterologous partners (WT *CEN1/CEN4*) (Fig. 2B). The mini-chromosome pair (WT *CEN^P^/CEN^P^*) shows higher variation than the homologous pair, but less than the independent movements of the heterologous partners (Fig. 2B). Notably, when we prevented centromere pairing in prophase (*zip1-N1*), the mini-chromosome centromeres were significantly less constrained (Fig. 2B), suggesting that centromere pairing in prophase connects the partner centromeres in prometaphase.

**Figure 2.**
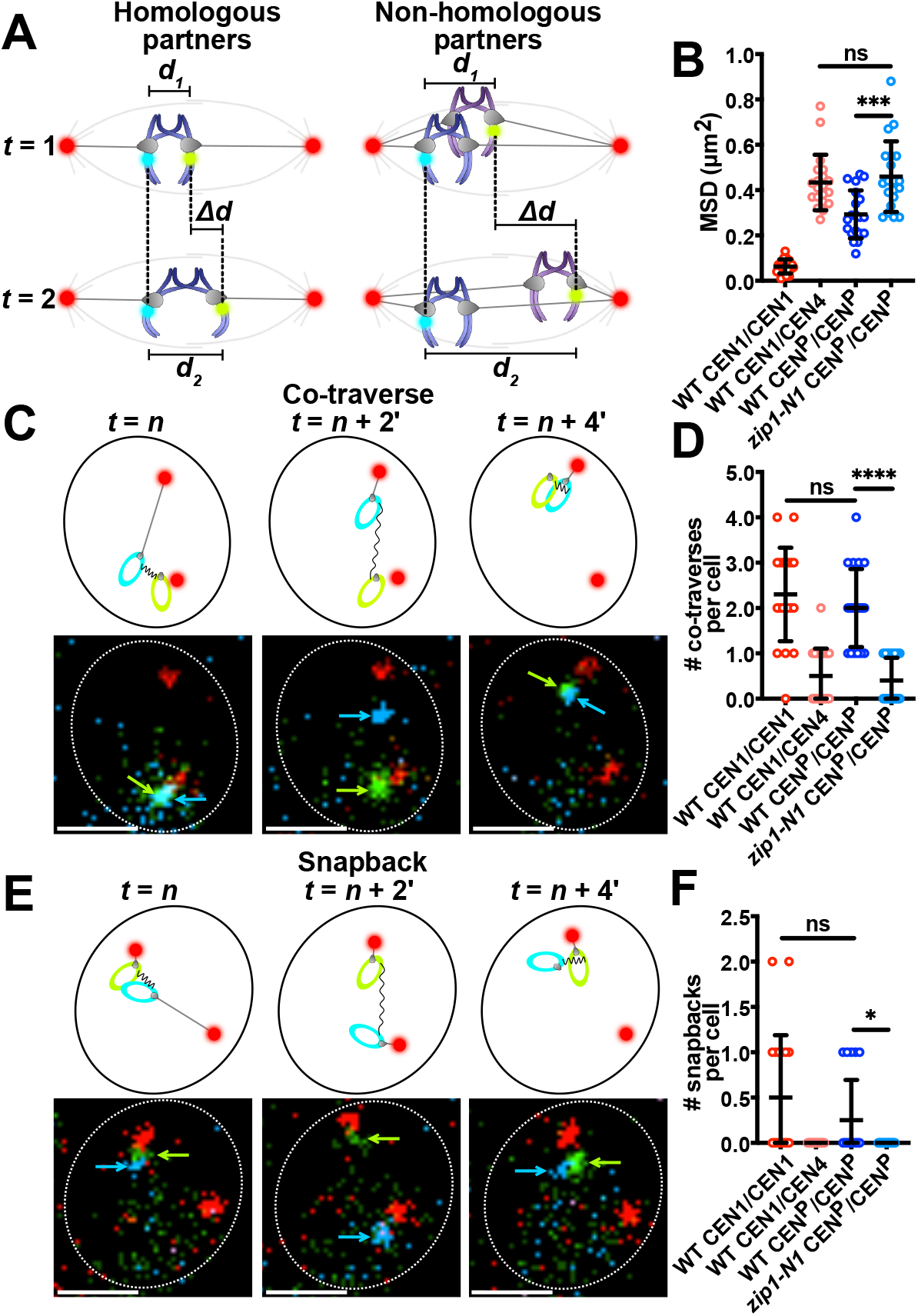
Partner centromeres exhibit behaviors consistent with inter-centromeric connections. **A)** Cartoon of methods used to measure inter-centromeric distances over time for homologous and non-homologous chromosomes. **B)** Mean-squared displacement (MSD) analysis was performed using twenty cells for each genotype. MSD was calculated by averaging the squares of the differences between each successive frame for the first ten frames (one every two minutes) after entry into prometaphase. **C)** Cartoons and representative micrographs showing a co-traverse. Arrows show locations of centromere plasmids. Scale bar = 2 μm. **D)** The number of co-traverses observed in the first ten frames after SPB separation in twenty cells per genotype. A co-traverse was scored when both the yEVenus and mTurquoise signals were within 0.5 μm of one SPB, then both moved to within 0.5 μm of the opposite SPB within three consecutive frames. A replicate of this experiment is shown in Supplemental Figure 2 A. **E)** Cartoons and representative micrographs depicting a snapback event. Arrows show relative locations of bi-orienting centromere plasmids. Scale bar = 2 μm. **F)** The number of snapbacks observed in the first ten frames after SPB separation in twenty cells per genotype. A snapback is defined as both yEVenus and mTurquoise signals being fully or partially colocalized (<0.3 μm separation), then separating to at least 0.6 μm, then re-colocalizing within three consecutive frames. A replicate of this experiment is shown in Supplemental Figure 2 B. Statistical comparisons were performed using unpaired t tests (*P<0.05, ***P<0.001, ****P<0.0001).

In meiosis I prometaphase, chromosomes track back and forth across the spindle until the homologous centromeres capture microtubules that radiate from opposite sides of the spindle^20^. Centromere partners that are tethered together due adjacent crossovers or a nascent inter-centromeric connection would be predicted to exhibit interdependence during these migrations. To test this, we defined two different behaviors as measures of interdependent movement and scored the frequency of these behaviors for the mini-chromosome pair and the control centromere pairs. First, we defined a “co-traverse” as both centromeres moving across the spindle within a four-minutes (a pole-to-pole traverse of the meiosis I spindle takes about two minutes^20^), with the expectation that if one centromere is pulled across the spindle by microtubule-dependent forces a connected partner centromere would sometimes be dragged with it (Fig. 2C). The incidence of co-traverses was significantly higher for a homologous centromere pair than for heterologous centromeres that show behave independently (Fig. 2D; *CEN1/CEN1* vs. *CEN1/CEN4*). This high frequency of co-traverses of the homologous centromeres was not because they individually exhibited more traverses than the heterologous centromeres - all centromeres assayed exhibited similar individual frequencies of moving across the spindle (Supl. Fig. 1 B). In wild-type cells the mini-chromosome pair exhibited similar frequencies of co-traverses as the homologous *CEN1/CEN1* centromeres (Fig. 2D; *CEN^P^/CEN^P^* vs. *CEN1/CEN1*). Notably, however, the mini-chromosome pair was more likely than the *CEN1/CEN1* pair (~30% vs 10% of co-traverses) to co-traverse separately, with one leading and the second following (Supl. Fig. 1 C), as if they were connected by a looser tether than the homologous centromeres. In the absence of centromere-pairing (*zip1-N1*) the mini-chromosome pair, like the heterologous control, showed no co-traverses.

As a second measure of interdependent movement, we defined a snapback as both centromeres being together, then one centromere moving away from then returning to its partner within a limited time frame (as if drawn back by an elastic connection) (Fig. 2E). This behavior was much more frequent in the movements of the homologous pair than the heterologous pair (Fig. 2F; *CEN1/CEN1* vs. *CEN1/CEN4*). In wildtype cells the minichromosomes mimicked the homologous pair (Fig. 2F; *CEN^P^/CEN^P^* vs. *CEN1/CEN1*). The snap-behavior was rare in *zip1-N1* strains without centromere-pairing (Fig. 2F).

## Centromere pairing promotes high fidelity meiosis I segregation

Previous studies using fixed-cell imaging have been unable to determine the true impact of centromere pairing on segregation fidelity. Centromere pairing of the various model chromosomes used in these studies occurs in only 50-70% of cells^4,5,16^ (as opposed to natural chromosome which pair their centromeres in virtually every meiosis). Previous measurements of meiotic segregation fidelity could not determine which model chromosomes had actually undergone pairing in prophase^10,12,21–23^. To address this, we used live cell imaging to identify mini-chromosomes that were either paired or not paired in prophase and then tracked their segregation fates in anaphase I. Centromeres were categorized as “paired” if they co-localized in three sequential frames of imaging in prophase prior to entry into prometaphase (SPB separation). Representative images of cells scored as paired (P) or unpaired (U) are shown in Fig. 3A and examples of cells scored as disjoined (DJ) or nondisjoined (NDJ) are shown in Fig. 3B. Consistent with prior fixed-cell studies, the mini-chromosomes exhibited about 75% correct segregation at meiosis I (without regard to prophase centromere pairing status) (Fig. 3C) and eliminating centromere pairing with a *zip1-N1* mutation randomized segregation of the minichromosomes^18^ (Fig. 3C).

**Figure 3.**
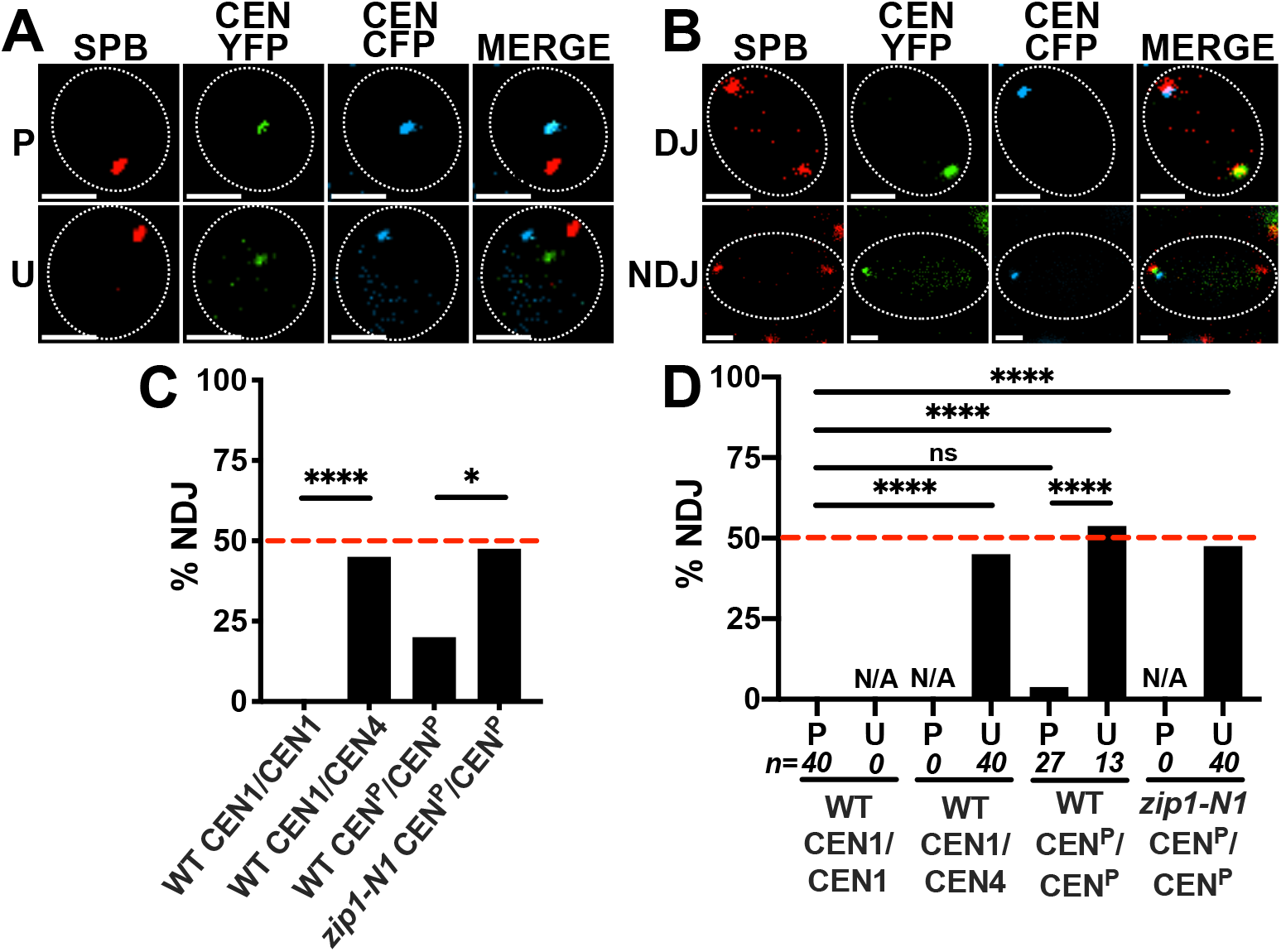
Centromere pairing in meiotic prophase ensures high fidelity segregation at anaphase I. **A)** Images were captured every 10 minutes. Representative micrographs showing examples of two cells in meiotic prophase (identified by a single SPB) in which the mTurquoise (blue) or yEVenus (green) tagged centromeres are either paired (P) or unpaired (U). Centromeres were classified as paired if they were colocalized (≤0.3 μm) for three consecutive frames (10 minutes apart) during prophase imaging. **B)** Representative micrographs showing two anaphase I cells (identified by two SPBs separated by at least 5 μm) in which the centromeres either disjoined (DJ) (segregated to opposite sides of the spindle) or nondisjoined (NDJ) (both centromeres on the same side of the spindle). **C)** Forty cells were scored for each genotype. **D)** Using the same forty cells from (**C**), we compared nondisjunction frequencies of each genotype in cells that were scored as having been paired (P) or unpaired (U) centromeres in prophase. Scale bars = 2 μm. Red dotted lines indicate random segregation. Statistical comparisons were performed using Fisher’s exact test (*P<0.05, ****P<0.0001).

In tracking individual chromosome pairs through meiosis I, we found that the centromeres of the homologous control chromosomes (*CEN1/CEN1*) were always paired just prior to prometaphase, and as expected, disjoined 100% of the time. The heterologous control centromeres (*CEN1/CEN4*) were never paired (Fig. 3D) and segregated randomly with respect to each other (each one presumably segregating from its own homologous partner). Consistent with previous studies mini-chromosomes were paired in ~65% of meiosis ^4,5,16^. Those that were paired exhibited over 95% correct segregation while those that were not paired segregated randomly (Fig. 3D).

The interdependent biorientation behaviors of the mini-chromosomes suggest that the centromeres of partner chromosomes become connected to one another during centromere pairing. These proposed connections could contribute to the bi-orientation process is by transmitting tension between the partner centromeres when they connect to microtubules that radiate from opposite sides of the spindle. Centromeres that are not under tension send a spindle checkpoint “wait” signal and non-exchange partner centromeres are able generate this signal^24,25^. If inter-centromeric connections that are formed during centromere-pairing can transmit tension between bi-oriented centromeres, this should satisfy the “wait” signal and allow cells to begin progressing to anaphase. We tested this prediction by monitoring yeast cells from prophase to anaphase and determining whether cells in which the mini-chromosomes disjoin exhibit a shorter prometaphase. First, to determine whether the centromere pairing process impacts bi-orientation times, we correlated the presence or absence of centromere pairing with the duration of prometaphase. In wild-type cells with either homologous (*CEN1/CEN1*) or heterologous (*CEN1/CEN4*) fluorescent centromere tags, cells spent an average of approximately eighty minutes in prometaphase (Fig. 4A). Cells carrying the mini-chromosomes spent about the same amount of time in prometaphase as the control strains if the minichromosomes had paired in prophase, but cells in which the mini-chromosomes had not paired spent significantly longer in prometaphase - an average of approximately 100 minutes. (Fig. 4A). When centromere pairing was abolished with the *zip1-N1* mutation (Fig. 4A; *zip1-N1 CEN^P^/CEN^P^*) the duration of prometaphase was also longer than in the control strains (Fig. 4B). Because the *zip1-N1* mutation randomizes disjunction for the mini-chromosome (Fig. 2D), we were able to determine whether cells in which the mini-chromosome disjoined as a result of random segregation were able to detect this correct outcome and exit prometaphase. Instead, the *zip1-N1* cells that disjoined or nondisjoined the mini-chromosomes exhibited indistinguishable prometaphase durations that were longer than those seen when the minichromosomes had undergone centromere pairing then disjoined. Thus, cells cannot detect correct segregation outcomes of non-connected partners and extinguish the spindle checkpoint “wait” signal.

**Figure 4.**
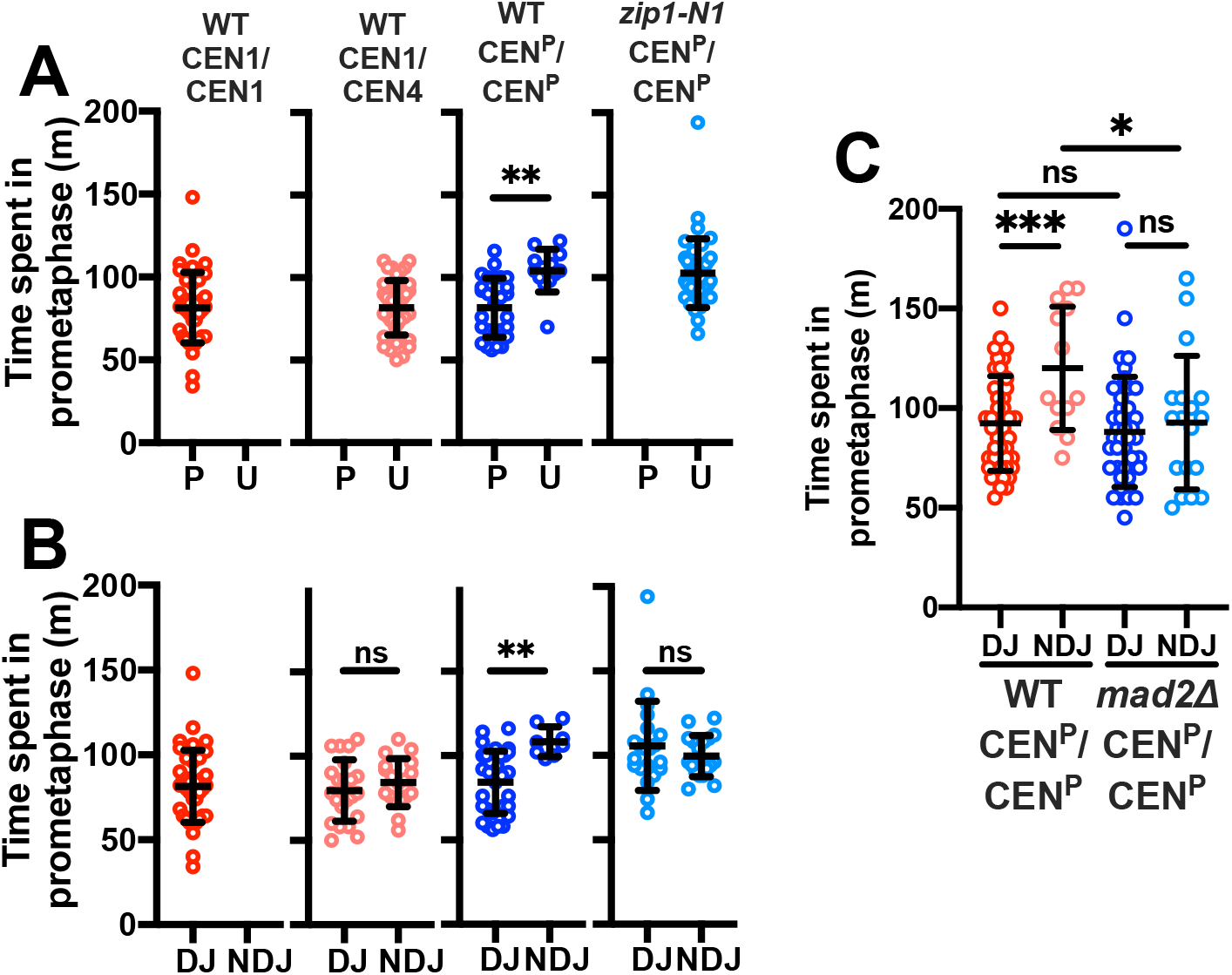
Inter-centromeric connections are sufficient to satisfy the spindle checkpoint in meiosis I. **A)** Forty prophase cells from each genotype were scored for whether their centromeres paired in prophase and their duration of prometaphase. **B)** The same forty cells from (**A**) were scored their segregation outcomes and their time spent in prometaphase. **C)** Wild-type cells and *mad2Δ* mutants were scored for segregation outcomes of the minichromosomes. The duration of prometaphase is graphed as a function of segregation outcome for each genotype. The combined data for three replicates of twenty cells are shown for each genotype. The nondisjunction frequencies for the *WT CEN^P^/CEN^P^* replicates were 6/20, 3/20, 4/20 and for the *mad2△ CEN^P^/CEN^P^* replicates were 7/20, 6/20, 5/20. Statistical comparisons were performed using unpaired *t* tests (*P<0.05, **P<0.01, ***P<0.001).

The shorter prometaphase durations of mini-chromosomes that disjoin following centromere-pairing (Fig. 4B; *CEN^P^/CEN^P^*) is consistent with the model that, like chiasmata, a bridge formed during centromere pairing can signal to the checkpoint machinery that biorientation has been achieved. To test this, we eliminated the spindle checkpoint (*mad2△*) and monitored metaphase duration for cells that did or did not correctly segregate the minichromosomes. Removing the checkpoint eliminated the delay seen in cells that nondisjoined the mini-chromosomes (Fig. 4C). Wild-type cells that disjoined the mini-chromosomes exhibited only slightly (but not significantly) longer metaphases than those that disjoined the minichromosomes in *mad2△* mutants. Thus, in the wild-type cells the mini-chromosomes that disjoin are not triggering a protracted delay beyond that triggered by the natural chromosomes. This suggests that chromosomes bi-orienting by virtue of centromere pairing alone do so just as quickly as the natural chromosomes, which we presume are linked at their centromeres and also by chiasmata. Further, since these cells transition with normal timing from metaphase to anaphase, the bi-oriented mini-chromosomes must effectively silence their spindle checkpoint “wait” signal – consistent with the transmission of tension between their bi-oriented kinetochores.

## Discussion

These analyses of bi-orientation behavior show that mini-chromosomes that have undergone centromere pairing are interdependent in their movements on the prometaphase spindle – suggesting they have become connected to one another. Inter-homolog chromatin threads have been observed previously between partner chromosomes in crane-fly spermatocytes^26^ and the small non-exchange chromosome 4 in Drosophila females^27^. Although we do not know the nature of these connections, one possibility is that during centromere-pairing, chromatin loops from the two partners become connected by cohesin, forming a flexible bridge that would explain the interdependent movements of the mini-chromosomes studies here. Indeed, we showed previously in budding yeast that prophase centromere pairing cannot ensure proper segregation in *sgo1△* mutants, in which cohesin is not protected at meiosis I centromere ^32^. Cohesin is enriched at meiotic centromeres and numerous studies suggest that cohesion establishment may be active in meiotic prophase^28–31^ (reviewed in ^32^) making cohesion a plausible candidate for linking partner centromeres.

The high fidelity of mini-chromosome segregation following centromere pairing has important implications for natural chromosomes. Although non-exchange partners, like the minichromosomes, have provided a unique way to observe how centromere pairing can influence disjunction, in budding yeast, non-exchange partners occur in only a few percent of meioses^33,34^. Our findings suggest that centromere pairing in meiotic prophase allows the formation of a physical connection between natural homologous chromosome partners, both those with and those without crossovers (Fig. 5). These connections could then contribute to biorientation in prometaphase either by providing a more direct connection between the kinetochores of exchange chromosomes as they bi-orient than is provided by more distal chiasmata, or by providing the sole connection between kinetochores of non-exchange pairs.

**Figure 5.**
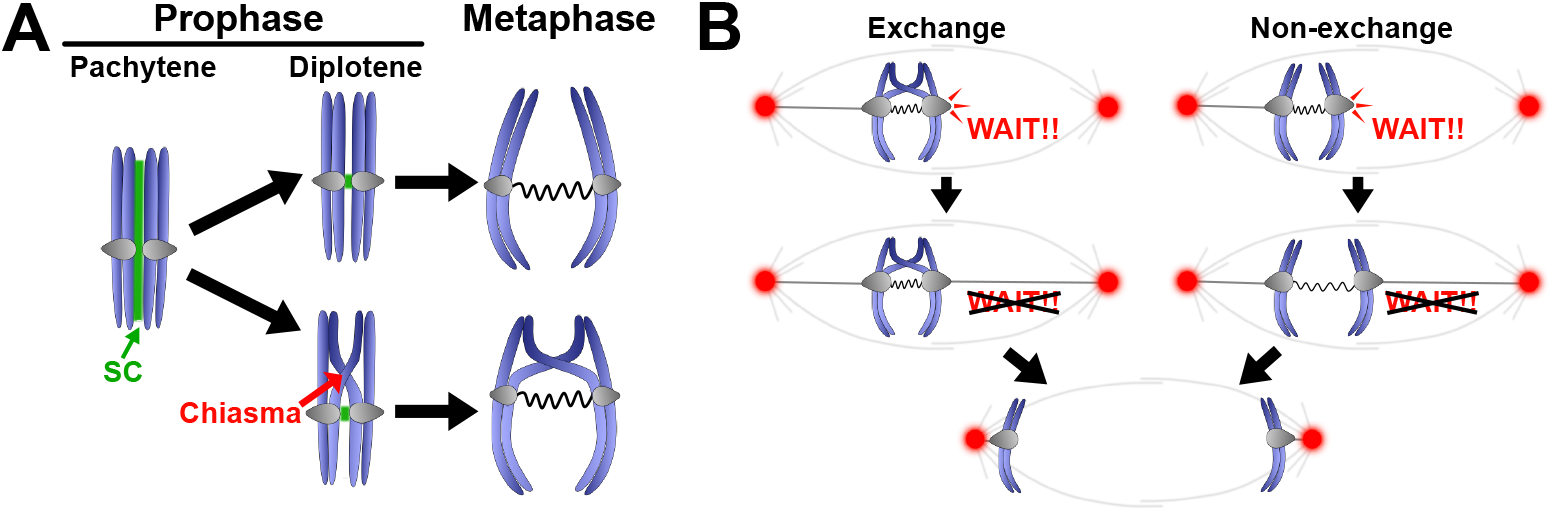
Model: Connections between homologous centromeres act with tension-dependent bi-orientation mechanisms to promote meiotic segregation. **A)** Crossing-over allows efficient synaptonemal complex (SC) assemble between homologous partners. After most of the SC disassembles, some SC components linger near the centromeres. The centromeres are “paired” in both exchange and non-exchange chromosomes. This pairing juxtaposes centromeric chromatin loops from homologous partners in the cohesin-rich centromeric regions. We propose this allows the linkage of chromatin from the partner centromeres by cohesin. **B)** Kinetochores that are unattached to the meiotic spindle emit a “WAIT!!” signal during biorientation. Bipolar attachments will generate tension across the bivalent, stabilizing the kinetochore-microtubule connection and turning off the “WAIT!!” signal. We propose that centromere pairing allows formation of a direct link between the bi-orienting meiotic centromeres of both exchange and non-exchange partners.

In yeast, fruit flies, and humans, exchanges are not equally able to insure disjunction and chromosomes with chiasmata far from the centromeres are more likely to nondisjoin^25,26^. Consistent with this, in budding yeast, the largest chromosomes, which often have a larger distance between the centromere and the closest chiasma, are especially dependent upon the spindle checkpoint for their segregation in meiosis^27^ and chromosomes with crossovers close to their centromeres bi-orient effectively, even without the spindle checkpoint^28^. These findings suggest that with increasing distance from the centromere, crossovers become decreasingly effective at transmitting signals between centromeres during the bi-orientation process. Given that the natural chromosome pairs in yeast and mouse spermatocytes (unlike the yeast model chromosomes) virtually all experience centromere pairing in meiotic prophase^11,15,29^, and the efficiency of correct segregation following centromere pairing, our data suggest that the biorientation of most natural chromosomes could be driven by direct centromere-to-centromere bridges formed during centromere pairing. By this model, crossing-over would have the critical role of driving the homologous synapsis that leads to centromere pairing and would serve as a second conduit over which tension could be transmitted during bi-orientation.

