## Supplementary material for "Centromere Pairing in Prophase Allows Partner Chromosomes to Orient on the Meiosis I Spindle": Methods, Supplementary Figures and Tables

**SUPPLEMENTAL MATERIALS**

**Methods**

**Yeast Strains and Growth**

The yeast strains used here are isogenic derivatives of rapidly sporulating strains of primarily SK1 and W303 ancestry, derived in the RE Esposito laboratory[1]. Complete sequences of the parental strains are available at NCBI. Diploid cells were grown overnight in YPAD medium or SD Sunrise -Leu, -Ura, -Trp medium for cells bearing mini-chromosome plasmids. To induce meiosis, overnight cultures were grown in YP acetate to 3-5x$10^7$ cells/mL and then shifted to 1% potassium acetate at $10^8$ cells/mL for five hours. Cells were then treated with 5 μM β-estradiol to synchronously release cells from pachytene arrest (see below).

**Fluorescence microscopy**

Live cell imaging was made possible using the CellASIC Onix Microfluidic system in conjunction with a Nikon Instruments Eclipse Ti2 inverted microscope. The microfluidic system allowed cells to be arranged in a monolayer while being continuously treated with fresh medium and, if required, reagents to induce cellular effects. The stage, objective lenses, and microfluidic plate were pre-warmed to 30°C before loading the cells. Individual chambers of the microfluidic plate were filled with fresh medium which is pumped into the cell chamber using pre-specified time intervals determined by the experimental protocol. Cells were loaded into the visualization chamber at a pressure of 8 psi for 20 seconds. The plate was placed on the microscope stage and an environmental chamber was placed over the plate, ensuring a constant 30°C temperature throughout the length of the experiment. Cells were induced to enter prometaphase by expression of the *NDT80* transcription factor. The addition of estradiol to the medium activates Gal4-ER, which turns on $P_{GAL1}$-NDT80[2,3]. Images were collected in five-slice Z-series in the mTurquoise and yEVenus channels to follow tagged centromeres and in the red channel for track the spindle pole marker Spc42-DSRed. The microscope (a Nikon Eclipse Ti2) was equipped with a Lumencor SPECTRA X light engine using an array of bandpass filters controlled by triggering system to minimize exposure times during image acquisition. A set of dichroic filter cubes was used to capture multiple channels per field per timepoint. Images were captured using a Hamamatsu ORCA-Flash4.0 digital CMOS camera. Nikon Instruments NIS Elements v5.0 was used to capture and save the time-lapse movies. Images were enhanced using NIS Elements software for the purposes of producing figures for publication. Distances were measured in three dimensions using Imaris software in the OMRF Imaging Core Facility

**Pairing, Movement, and Disjunction Analyses**

Immediately after initiating the release from pachytene (addition of estradiol), cells were imaged every ten minutes for one hour. This low-frequency imaging timeframe was sufficient to identify cells with paired or unpaired chromosomes, yet infrequent enough to reduce phototoxic effects of imaging during prophase. Next, as cells began to enter prometaphase, we imaged every two minutes for three hours. Two minutes was chosen to reduce photobleaching over time while still revealing traverses, as described previously[4]. Lastly, as cells were exiting meiosis I we imaged every ten minutes for another hour to capture anaphase I cells to score for disjunction. Only the cells that satisfied all four of the following requirements were scored: 1) had sufficient fluorescent signals throughout meiosis to continuously track the mTurquoise and yEVenus tagged centromeres, 2) were able to be scored as paired or unpaired in prophase (Centromeres were classified as paired if they were colocalized (≤0.3 μm) for three consecutive frames (10 minutes apart) during prophase imaging.), 3) completed anaphase I, and 4) contained one copy of each of the two tagged centromeres. The mini-chromosomes are lost mitotically in a small fraction of cell divisions. Meiotic cells typically carried one or zero copies of each mini-chromosome. When only scoring for disjunction was necessary (Fig. 4C), cells were imaged starting one hour after release from pachytene arrest every ten minutes for four hours to capture anaphase I cells.

### Statistical Tests

Statistical tests were performed using Prism software.

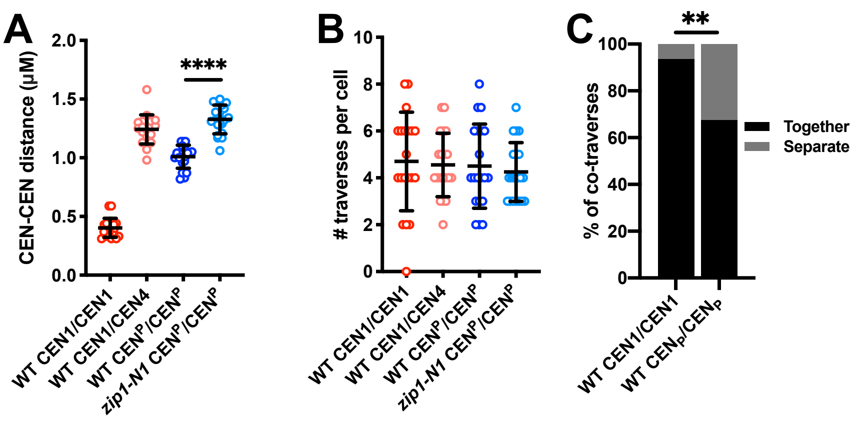


**Supplemental Figure 1: Monitoring partner centromere movements on the meiosis I spindle. A)** In the experiment described in Figure 2, inter-centromeric distances (CEN-CEN distance) were recorded for the first ten frames (one frame every two minutes) following entry into prometaphase (spindle pole body separation). The average distance between the two centromeres of each cell over this ten-minute interval is plotted in the figure. The average distance between the mini-chromosome centromeres is significantly larger if they have not paired prior to prometaphase entry. Statistical comparisons were performed using an unpaired *t* test (****P<0.0001). **B)** In the experiment shown in Figure 2, we scored the combined number of traverses exhibited by the two tagged centromeres in each cell over the first twenty minutes following entry into prometaphase (one frame every two minutes). A centromere was scored as having exhibited an individual traverse when it moved from a position within 0.5 μm of one SPB, then moved across the spindle to a position within 0.5 μm of the opposite SPB within three consecutive frames. Although the frequency of co-traverses (both centromeres making the same traverse in the same time window) differed significantly between strains (Fig.2) there was no difference in the number of individual traverses exhibited by the tagged centromeres in these tie frames. **C)** Co-traverses (Fig. 2) occurred when both tagged centromeres in a strain crossed the spindle within the same four-minute time interval.  These co-traverses could be classified into two categories. Co-traverses classified as “separate” fulfilled the criteria of a co-traverse as defined in Figure 2 but the yEVenus and mTurquoise centromere tags were not touching. Co-traverses classified as “together” when the centromere tags were close enough together that they were overlapping. Statistical comparisons of relevant groups were performed using Fisher’s exact test. (**$P<0.01$). Although for both the mini-chromosomes and chromosome *I*, the centromere tags were usually over-lapping, the mini-chromosomes had a significantly higher fraction that co-traversed as separate yEVenus and mTurquoise signals.

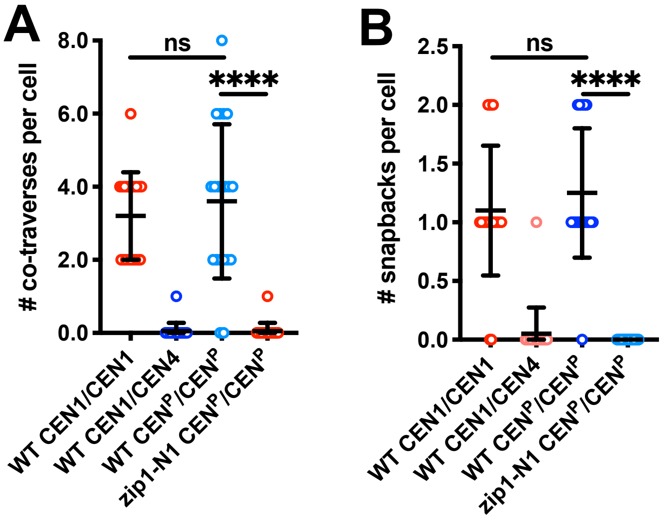


**Supplemental Figure 2: Independent replicates of the co-traverse and snapback experiments.** Independent biological replicates were performed for the co-traverse and snapback experiments shown in Figure 1. **A)** The number of co-traverses observed in the first ten frames (one every two minutes) after entry into prometaphase (spindle pole body separation) in twenty cells per genotype. A co-traverse is defined (as in Fig. 2 C) as both the yEVenus and mTurquoise centromere tag signals being positioned within 0.5 µm of one SPB, then both moving to within 0.5 µm of the opposite SPB within three consecutive frames (four minutes). **B)** The number of snapbacks observed in the first ten frames after entry into prometaphase (SPB separation) in twenty cells per genotype. A snapback is defined as both the yEVenus and mTurquoise centromere tag signals being fully or partially colocalized (<0.3 µm), then separating to at least 0.6 µm, then re-colocalizing within three consecutive frames (as in Fig. 2 E). Statistical comparisons were performed using an unpaired *t* test (****P<0.0001). The results of both replicates mirror those of the version shown in Figure 2.

**Table S1: Diploid Strain List**

| Diploid | Parent strains | Identifier | Figure(s) |
|---|---|---|---|
| DJE115 | X3661 x Y3001 | WT CEN1/CEN1 | Fig. 2B, 2D, 2F, 3C, 3D, 4A, 44B |
| DJE117 | X3660 x Y3065 | WT CEN1/CEN4 | Fig. 2B, 2D, 2F, 3C, 3D, 4A, 4B |
| DJE118 | X3101 x Y2735 | WT $CEN^P/CEN^P$ | Fig. 1, 2B, 2D, 2F, 3C, 3D, 4A, 4B, 4C |
| DJE129 | X3706 x Y3042 | *zip1-N1* $CEN^P/CEN^P$ | Fig. 2B, 2D, 2F, 3C, 3D, 4A, 4B |
| DJE161 | X3766 x Y3113 | *mad2Δ* $CEN^P/CEN^P$ | Fig. 4C |

**Table S2: Haploid Strain List**

| | |
|---|---|
| X3661 | *MATa, ura3-13, trp1-Δ63, leu2-?, tyr1-1, lys2-1, can1-R, his3::pHC30[pDMC1-3x yEVenus-lacI-I12 HIS3], SPC42-[MDE1145: URA3::HIS SPC42-DSRed], ura3::pKB80 [PGPD1-GAL4(848)-ER-URA3::hphNT1], natNT2-PGAL1-NDT80, CEN1::pJN2[lacO256 LEU2]* |
| Y3001 | *MATα, leu2-?, lys2-2, met13-c, tyr1-2, ura3-1, trp1-Δ63, cyh2-1, his3::pHC29[pURA3-tetR-3XmTurquoise-9MYC HIS3], trp1::pD280[pDMC1-3XyEVenus-lacI-I12 TRP1], natNT2-PGAL1-NDT80, CEN1::pMNS18[tetO256 URA3]* |
| X3660 | *MATα, ura3-13, trp1-Δ63, leu2-?, tyr1-1, lys2-1, can1-R, his3::pHC30[pDMC1-3x yEVenus-lacI-I12 HIS3], SPC42-[MDE1145: URA3::HIS SPC42-DSRed], ura3::pKB80 [PGPD1-GAL4(848)-ER-URA3::hphNT1], natNT2-PGAL1-NDT80, CEN1::pJN2[lacO256 LEU2]* |
| Y3065 | *MATa, leu2-?, lys2-2, met13-c, tyr1-2, ura3-1, trp1-Δ63, cyh2-1, his3::pHC29[pURA3-tetR-3XmTurquoise-9MYC HIS3], trp1::pD280[pDMC1-3XyEVenus-lacI-I12 TRP1], natNT2-PGAL1-NDT80, CEN4::pMNS23[tetO256 URA3]* |
| X3101 | *MATa, ura3-13, trp1-Δ63, leu2-?, tyr1-1, lys2-1, can1-R, his3::pHC30[pDMC1-3x yEVenus-lacI-I12 HIS3], SPC42-[MDE1145: URA3::HIS SPC42-DSRed], ura3::pKB80 [PGPD1-GAL4(848)-ER-URA3::hphNT1], natNT2-PGAL1-NDT80, OPL210 [lacO, CEN3, TRP1/ARS1, LEU2]* |
| Y2735 | *MATα, leu2-?, lys2-2, met13-c, tyr1-2, ura3-1, trp1-Δ63, cyh2-1, his3::pHC29[pURA3-tetR-3XmTurquoise-9MYC HIS3], trp1::pD280[pDMC1-3XyEVenus-lacI-I12 TRP1], natNT2-PGAL1-NDT80, OPL214 [tetO, CEN3, TRP1/ARS1, URA3]* |
| X3706 | *MATα, ura3-13, trp1-Δ63, leu2-?, tyr1-1, lys2-1, can1-R, his3::pHC30[pDMC1-3x yEVenus-lacI-I12 HIS3], SPC42-[MDE1145: URA3::HIS SPC42-DSRed], ura3::pKB80 [PGPD1-GAL4(848)-ER-URA3::hphNT1], natNT2-PGAL1-NDT80, zip1::pELK11[zip1-N1]-kanMX, OPL210 [lacO, CEN3, TRP1, LEU2]* |
| Y3042 | *MATa, leu2-?, lys2-2, met13-c, tyr1-2, ura3-1, trp1-Δ63, cyh2-1, his3::pHC29[pURA3-tetR-3XmTurquoise-9MYC HIS3], trp1::pD280[pDMC1-3XyEVenus-lacI-I12 TRP1], natNT2-PGAL1-NDT80, zip1::pELK11[zip1-N1]-kanMX, OPL214 [tetO, CEN3, TRP1/ARS1, URA3]* |
| X3766 | *MATa, ura3-13, trp1-Δ63, leu2-?, tyr1-1, lys2-1, can1-R, his3::pHC30[pDMC1-3x yEVenus-lacI-I12 HIS3], SPC42-[MDE1145: URA3::HIS SPC42-DSRed], ura3::pKB80 [PGPD1-GAL4(848)-ER-URA3::hphNT1], natNT2-PGAL1-NDT80, mad2::NATMX6, OPL210 [lacO, CEN3, TRP1/ARS1, LEU2]* |
| Y3113 | *MATα, leu2-?, lys2-2, met13-c, tyr1-2, ura3-1, trp1-Δ63, cyh2-1, his3::pHC29[pURA3-tetR-3XmTurquoise-9MYC HIS3], trp1::pD280[pDMC1-3XyEVenus-lacI-I12 TRP1], natNT2-PGAL1-NDT80, mad2::NATMX6, OPL214 [tetO, CEN3, TRP1/ARS1, URA3]* |
